# Type II muscle fibre properties are not associated with balance recovery following large perturbations during walking in young and older adults

**DOI:** 10.1101/2021.11.26.470167

**Authors:** Christopher McCrum, Lotte Grevendonk, Gert Schaart, Esther Moonen-Kornips, Johanna. A. Jörgensen, Anne Gemmink, Kenneth Meijer, Joris Hoeks

## Abstract

Falls among older adults are often attributed to declining muscle strength with ageing. Associations between muscle strength and balance control have been reported, but the evidence for, and key mechanisms of resistance exercise in fall prevention are unclear. No studies have directly examined the relationship between muscle fibre characteristics and reactive balance control. Here, we address whether or not Type II muscle fibre characteristics associate with reactive balance during walking in young and older adults with varying muscle fibre type composition. We analyse muscle biopsy-derived fibre characteristics and stability during a treadmill-based walking perturbation (trip-like) task of healthy young adults, healthy, normally active older adults, trained older adults and physically impaired older adults. We find no significant associations between Type II muscle fibre properties and reactive balance during walking, indicating that practitioners and researchers should consider more than just the muscle tissue properties when assessing and intervening on fall risk.

## Introduction

Loss of muscle mass in humans after the age of 50 years is largely attributable to declines in lower body muscle mass [1], which has implications for mobility in older age. Specifically, the leg extensor muscle-tendon groups critical for gait [2–4] show reduced muscle strength and altered tendon mechanical properties in older, compared to younger adults [5–8]. Lower limb muscle strength [9–12], power [11,13,14] and quality [15,16] have been associated with daily life falls incidence in older adults, although findings are not unequivocal [12,13,17–19]. The majority of falls in community-dwelling older adults occur following large balance disturbances (like trips and slips) during walking [20–27]. Small to moderate associations have been reported between lower limb muscle strength and balance recovery performance following laboratory-based lean-and-release [28–32], trip [33–36] and slip [37,38] perturbations simulating common causes of falls. Regarding interventions, there is weak or mixed evidence that general strength training alone can improve balance [39–41] and reduce falls [42,43], and whether a direct cause-effect relationship between muscle strength and balance performance exists has been questioned [44].

The loss in muscle strength and function with increasing age may occur due to muscle fibre atrophy and muscle fibre loss [45–48]. Age-related decline in muscle mass seems to be primarily attributable to an atrophy of specifically Type II muscle fibres [49–54]. Older age is also associated with a change in the expression of myosin heavy chain isoforms (MHC) in favor of slow MHC I, reflecting the shift towards a slower fibre type profile (Type I) and the selective atrophy of Type II fibres [54–56]. One study on older female participants with and without hip fractures reported that age and hip fracture history had significant individual and accumulating effects on Type II muscle fibre size (decreasing with age and fall-related hip fracture) [49]. Encouragingly, physical activity may prevent or slow this age-related atrophy of Type II fibres [52] and it has been reported that increases in muscle mass as a result of resistance exercise in old age is attributable to hypertrophy of Type II fibres specifically (and almost exclusively)[51]. In contrast, in endurance-trained older adults, greater proportions of Type I fibres have been observed when compared to strength-trained older adults, untrained older adults and untrained young individuals [57–59].

Together, these data suggest that muscle fibre characteristics may be an important factor in balance performance and falls. Despite this, there are no studies, to our knowledge, directly examining the relationship between muscle fibre characteristics and reactive balance control in the literature. In this study, we address for the first time whether or not Type II muscle fibre characteristics associate with reactive balance during walking in young and older adults with varying muscle fibre type composition. To do this, we analyse muscle biopsy-derived fibre characteristics and stability during a treadmill-based gait perturbation task of healthy young adults, healthy, normally active older adults, trained older adults and physically impaired older adults. Based on previous studies, we hypothesised that there would be a small to moderate relationship between Type II muscle fibre characteristics (size and proportion) and stability recovery following large perturbations during walking (number of recovery steps needed following a novel perturbation). We additionally explored these relationships after repeated perturbations to examine the potential role of Type II muscle fibre characteristics in adaptability of stability recovery responses.

## Methods

### Participants

The data and analyses in the current secondary analysis were part of a larger study (n=59), which was approved by the institutional Medical Ethical Committee, conducted in accordance with the declaration of Helsinki and registered at clinicaltrials.gov (NCT03666013). All participants provided their written informed consent. Some data have been reported in our previous publication addressing different research questions [60]. For the current secondary analysis, the data of all participants who underwent muscle biopsies and completed the walking perturbation experiment in the larger study were included. This resulted in a sample of 52 participants including 14 young (7 male and 7 female) and 38 older (22 male and 16 female) individuals. Due to this being a secondary analysis, no formal power calculation was conducted to determine the sample size. However, sample sizes of 46 and 47 will achieve approximately 80% power with a significance level of 0.05 to detect Pearson and Spearman Correlation Coefficients of 0.4, which we consider reasonable for the purpose of the current analysis.

Prior to inclusion, all participants underwent a medical screening that included a medical questionnaire, a physical examination by a physician, and an assessment of physical function by means of a Short Physical Performance Battery [61]. After screening, participants were assigned to the following groups: Young individuals with normal physical activity (Y, 20 – 30 years), older adults with normal physical activity (O, 65 – 80 years), trained older adults (TO, 65 – 80 years) and older adults with impaired physical function (IO, 65 – 80 years). Participants were considered normally physically active if they completed no more than one structured exercise session per week, whereas participants were considered trained if they engaged in at least 3 structured exercise sessions of at least 1 hour each per week for an uninterrupted period of at least one year. Participants were classified as older adults with impaired physical function (IO) in case of an SPPB score of ≤ 9.

### Body Composition and Muscle Volume

Body composition (fat and fat-free mass) was determined at 8 AM after an overnight fast from 10 PM the previous evening using air displacement plethysmography (BodPod®, COSMED, Inc., Rome, Italy) [62]. To measure muscle volume, a 3T whole body MRI scanner was used (Achieva 3T-X; Philips Healthcare, Best, The Netherlands). Participants were positioned in the supine position in the MR scanner (feet first) to determine muscle volume of the upper leg using the body coil. Subsequently, a series of T1-weighted images were acquired of the upper leg (slice thickness of 10mm, no gap between slices, in-plane resolution of 0.78 × 0.78mm). A custom-written MATLAB script was used to automatically segment adipose and muscle tissue. Thereby each image was normalised, and a histogram calculated. Histogram peaks corresponding to muscle tissue grey values were summed up, resulting in the count of muscle pixels. Muscle volume was calculated by voxel size times the number of muscle pixels: Vol_muscle =Vox_size*N_muscle pixels. Every slice was reviewed individually and in case of a clear notable deviation, a manual threshold correction for muscle grey values was performed. The muscle segmentation was performed in the consecutive slices between the starting point (tendon attachment) of the *m. rectus femoris* and of the *m. gluteus maximus*.

### Habitual physical activity

Habitual physical activity was estimated using an ActivPAL activity monitor (PAL Technologies, Glasgow, Scotland) for five consecutive days, including two weekend days. The total amount of steps per day was measured, as well as the total stepping time in proportion to waking time, determined according to van der Berg et al. [63]. Stepping time (i. e., physical activity) was then further classified into high-intensity physical activity (HPA; minutes with a step frequency >110 steps/min in proportion to waking time) [64].

### Muscle Strength

Maximum voluntary knee extension and flexion torque was measured using the Biodex System 3 Pro dynamometer (Biodex® Medical Systems, Inc., Shirley, NY, USA). Participants were stabilized with shoulder, leg, and abdominal straps to prevent compensatory movement and the test was performed with the left leg in all participants. Participants performed three 5s maximal extensions and flexions with a 30s rest period in between trials. The knee position was fixed at a 70° angle. Maximal voluntary isometric knee-extensor and knee-flexor torque was defined as the average of the highest two out of three peak torques. Participants also performed 30 consecutive extension and flexion movements (range of motion 120 degrees/s). The peak torque of each extension and flexion was recorded and the maxima were defined as the highest peak. For both protocols, maximal torque was recorded in Newton-meters and corrected for bodyweight (Nm/kg).

### Muscle Biopsy and Fibre Typing

A biopsy specimen was taken from the vastus lateralis muscle under local anesthesia (1.0% lidocaine without epinephrine), according to the Bergström method [65]. Muscle tissue was immediately frozen in melting isopentane cooled with liquid nitrogen for immunohistochemistry and stored at −80°C degrees until analysis. Fresh 5μm-thick cryosections were treated for five minutes with 0.25% Triton X-100 in phosphate-buffered saline and incubated for 45 minutes at room temperature with a mouse monoclonal MHC1-antibody (A.4840; Developmental Studies Hybridoma Bank, Iowa City, IO) and with the basal membrane marker laminin (L9393; Sigma, Zwijndrecht, The Netherlands). Thereafter, sections were incubated for 45 minutes with the appropriate secondary antibodies goat anti-mouse IgM AlexaFluor488 (A21042; Invitrogen Thermo Fisher, Breda, The Netherlands) and goat anti-rabbit IgG AlexaFluor555 (A21428; Invitrogen Thermo Fisher) and mounted with Mowiol. All sections were captured using a Nikon E800 fluorescence microscope (Nikon Instruments, Amsterdam, The Netherlands [66]) and images were analyzed in ImageJ [67]. Cell membranes were background corrected and threshold was set manually to create a binary image. Subsequently, the original cell membrane image (laminin staining) was added to the binary image of the cell membranes to check whether all cell membranes in the binary image were closed. If not, cell membranes were closed manually. Fibre size was determined by measuring the cross-sectional area (CSA) and fibre type distribution was expressed as a percentage with respect to the number of fibres and as a percentage of CSA (% fibre type I/II area relative to total CSA).

### Stability during Perturbed Walking

The walking measurements were conducted using the Computer Assisted Rehabilitation Environment Extended (CAREN; Motekforce Link, Amsterdam). The system includes a dual-belt force plate-instrumented treadmill (1000Hz), a 12 camera Vicon Nexus motion capture system (100Hz; Vicon Motion Systems, Oxford, UK) and a 180° virtual environment providing optic flow. A safety harness connected to an overhead frame was worn by participants at all times. Six retroflective markers were attached to anatomical landmarks (C7, sacrum, left and right trochanter and left and right hallux).

Participants were already accustomed to the CAREN setup, as they had previously completed an instrumented six-minute walk test as part of the larger study. To further ensure participants were comfortable and familiarised to the measurement setup and conditions, the perturbation measurement session began with unrecorded fixed-speed walking trials of two minutes at speeds of 0.4m/s up to 1.8m/s in 0.2m/s increments. Following a break, the same fixed-speed walking trials were repeated and recorded. During another break, the data from these trials were used to calculate a stability-normalised walking speed for each participant individually for use in the subsequent walking perturbation trial, which was set for a margin of stability (MoS; see below; [68]) of 0.05m as described previously [69], ensuring a comparable baseline walking stability for all groups [70–73]. Next, participants completed the perturbation trial which started with three to four minutes of unperturbed walking at the stability-normalised walking speed, followed by 10 unannounced unilateral treadmill belt acceleration perturbations (every 30 to 90 seconds). The first and tenth accelerations were applied to the right leg, while the second to ninth accelerations were applied to the left leg as described previously [70,71]. Participants were told that they would complete a walking balance challenge and to try to continue walking as normally as possible. The participants were unaware of the specifics of the perturbations and no warnings or cues were given during the trial. For the current study, the first perturbations to each limb were analysed (Pert1_R_ and Pert2_L_), representing novel disturbances, as well as the ninth perturbation (final left leg perturbation; Pert9_L_) to indicate adaptation in gait following eight repeated perturbations.

Data processing was conducted in MATLAB (2016a, The MathWorks, Inc., Natick). The three-dimensional coordinates of the markers were filtered using a low pass second order Butterworth filter (zero-phase) with a 12Hz cut-off frequency. Foot touchdown and toe-off were detected using a combined method of force plate data (50 N threshold) and foot marker data [74], as described in detail previously [75]. The anteroposterior MoS at foot touchdown were calculated as the anteroposterior distance between the anterior boundary of the base of support (BoS) and the extrapolated centre of mass (X_CoM_) [68], adapted for our validated reduced kinematic model [76] and treadmill walking [77], in the same manner as our previous studies [69–72]. The MoS was calculated for the following steps: the mean MoS of the eleventh to second last step before each perturbation (Base); the final step before each perturbation (Pre); and the first eight recovery steps following each perturbation (Post1-8). In order to determine the number of steps to recover to baseline stability (up to the eighth recovery step), two methods were used. The first, as used in our previous studies [71–73], used a 0.05m threshold – if the recovery step MoS value was within 0.05m of the Base MoS, this was heuristically considered not meaningfully different to unperturbed baseline walking. The second method, rather than using a standard value for all participants, used an individual threshold of three standard deviations of the 10 pre-perturbation steps used to calculate Base. This accounted for those individuals with either much lower or higher variability in their unperturbed walking for whom the 0.05m threshold might have been inappropriate. As there is no accepted standard for establishing the number of recovery steps, we have included and analysed both outcomes.

### Statistics

Normality for all data was checked with Shapiro-Wilk tests, which determined whether parametric or non-parametric analyses would be used. Group differences in the parameters were assessed using either one-way ANOVAs with Tukey’s multiple comparisons tests or Kruskal-Wallis tests with Dunn’s multiple comparisons tests. All analyses involving the number of recovery steps included analyses using the standard 0.05m threshold (Standard Threshold) and the individualised, three standard deviation threshold (Individual Threshold). Spearman correlations were conducted to assess the relationships between the recovery steps required by participants during Pert1_R_, Pert2_L_ and Pert9_L_ and the muscle fibre properties (percentage of Type II fibres, mean Type II fibre CSA and the percentage of CSA taken up by Type II fibres). Spearman correlations were also conducted for the recovery steps required and the muscle volume and strength measures. Correlations with Pert1_R_ and Pert2_L_ were used to determine associations with balance recovery following a novel perturbation (similar unexpected and unfamiliar situation to how falls occur in daily life) whereas associations with Pert9_L_ were used to determine associations with adaptability of the stability recovery responses. α was set at 0.05. Correlation coefficients were interpreted as follows: 0-0.1: Negligible; 0.1-0.3: Small; 0.3-0.5: Moderate; 0.5 and above: Large [78], although these may be overestimates in comparison to the field of gerontology, where there is evidence that Pearson’s r = 0.12, 0.20, and .032 may better represent small medium and large effects [79]. Statistical analyses were conducted, and figures were prepared using GraphPad Prism version 9 for Windows (GraphPad Software, LLC, San Diego, California, USA).

## Results

### Participant Characteristics

Of the 52 participants who were included in this study, one participant was excluded due to problems in both walking motion capture (noise and marker gaps) and muscle biopsy (not enough tissue), and a further three participants were excluded due to issues with the muscle biopsy (not enough useable tissue was collected). The remaining 48 participants’ data were included in the analyses and their characteristics, arranged per group, are displayed in Table 1. Note that we did not specifically aim to have groups of a specific size, as the groups themselves were not related to our primary aims. Rather we wanted to achieve a sample that included a wide range of muscle fibre type characteristics and stability recovery performance that could provide insight into the associations between these parameters. Due to technical issues, walking data from Pert1_R_ for one participant of group O, walking data from Pert2_L_ for one participant of group TO and MRI-derived muscle volume data for one participant of group TO were excluded.

**Table 1:** Participant characteristics, presented as mean (SD).

|  | Young, normal physical activity (Y) | Older adults, normal physical activity (O) | Physically impaired older adults (IO) | Trained older adults (TO) |
| --- | --- | --- | --- | --- |
| **N** | 12 | 16 | 3 | 17 |
| **Male** | 7 | 8 | 2 | 11 |
| **Age [years]** | 23.9 (2.2) | 70.6 (3) | 69.7 (4.2) | 67.8 (2.4) |
| **BMI [kg/m<sup>2</sup>]** | 22.4 (3.3) | 25.8 (3.4) | 28.4 (1.8) | 23.8 (1.8) |
| **Height [cm]** | 176 (7.4) | 169 (11.3) | 171 (4.4) | 171.2 (7.2) |
| **Weight [kg]** | 69.3 (11.5) | 74.0 (12.3) | 82.9 (2.4) | 70 (7.5) |
| **Fat mass [%]** | 22.6 (8.4) | 33.3 (9.5) | 37 (10.7) | 25.9 (8.1) |
| **SPPB 4m walk speed [m/s]** | 1.1 (0.2) | 1.2 (0.2) | 1.0 (0.3) | 1.3 (0.2) |
| **SPPB Chair-stand test [s]** | 9.2 (2.7) | 10.2 (1.4) | 13.7 (7.1) | 8.9 (2.1) |
| **Steps/day** | 10487 (3134) | 10007 (2885)‡ | 7594 (1851) | 13604 (6218) |
| **HPA time/wake time (%)** | 2.6 (1.7)* | 2.1 (1.2)‡ | 1.6 (0.1) | 4.9 (3.9) |

BMI: body mass index. SPPB: short physical performance battery. HPA: high intensity physical activity. \*: One participant excluded due to data processing issues. ‡: Two participants excluded due to data collection and analysis issues.

Groups O and Y possessed similar physical activity levels (~10K steps/day and similar active time in high-intensity activities; Table 1). Group TO, in comparison were more active (~13K steps/day and 4.9% active time was at high intensity; Table 1) and were mainly involved in endurance sports and training, whereas Group IO were less active (~7.5K steps/day and 1.6% active time at high intensity; Table 1).

### Type I and II Muscle Fibre Characteristics

In order to evaluate the muscle fibre type characteristics of the muscle biopsy samples, a mean (SD) of 454(260) muscle fibres were analysed per participant. Representative fluorescence microscope images used for fibre type analyses are displayed in Fig. 1. In order to explore the range in the characteristics, one-way ANOVAs were applied and did not reveal a significant group effect on Type I or II muscle fibre percentage (F(3, 44)=2.8, P=0.051), Type I fibre CSA (F(3, 44)=0.8, P=0.5) or Type II fibre CSA (F(3, 44)=1.9, P=0.14), although group TO tended to have lower Type II fibre percentage (Fig. 1). A one-way ANOVA did reveal a significant effect of group on the percentage of total CSA taken up by Type I and II fibres (F(3, 44)=4.5, P=0.007), with Tukey’s multiple comparisons tests revealing significantly lower values for group TO vs. Y (adjusted P=0.008) and O (adjusted P=0.036), as indicated in Fig. 1. While group comparisons were not always significant, these data confirm a range of muscle fibre type characteristics were present in our analysed participants, that there were no significant differences between groups Y and O and that group TO have an altered fibre type composition favouring Type I fibres. As can be seen in Fig. 1, Type II fibre mean CSA appeared to be lower in all older adult groups compared to group Y, despite a non-significant one-way ANOVA. *Post hoc* Mann Whitney tests comparing group Y (n=12, Median=5477μm^2^) with all older participants collapsed into one group (n=36, Median=3509μm^2^) and with Group O (n=16, Median=3500 μm^2^) revealed significant differences in Type II fibre mean CSA (U=114, P=0.014 and U=49, P=0.029, respectively).

**Figure 1:**
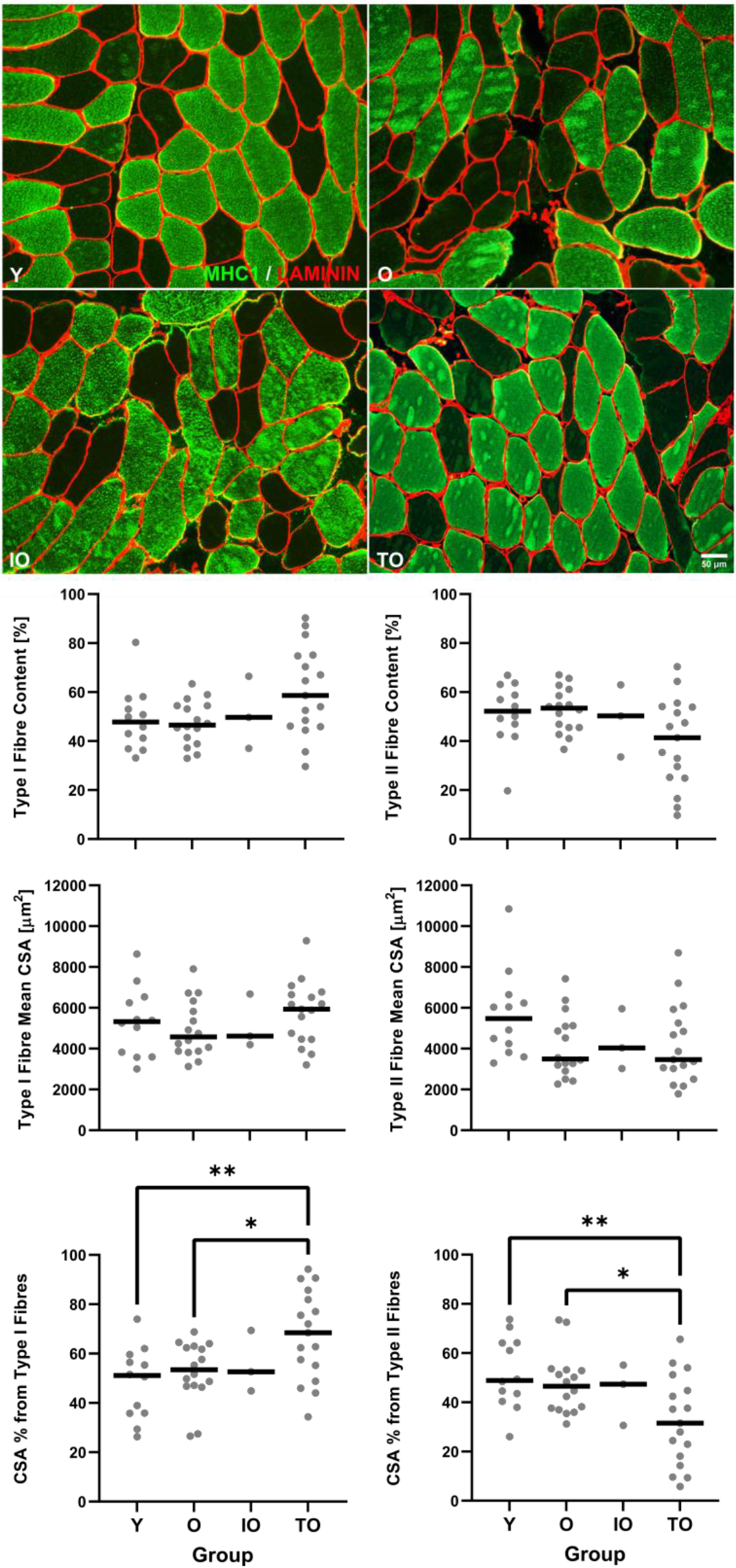
**Top Panel:** Representative fluorescence microscope images used for fibre type analyses from young adults (Y), older adults with normal physical activity (O), trained older adults (TO) and physically impaired older adults (IO). Basal membrane (laminin; red); type 1 fibres (MHC1; green) and type 2 fibres (no stain / black). Each participant shown has the median value of average fibre CSA within their group. **Bottom Panel:** Group median and individual Type I and II muscle fibre characteristics. *: P=0.036 **: P=0.008

### Muscle Volume, Strength Characteristics and Perturbation Recovery Steps

A one-way ANOVA did not reveal a significant effect of group on upper leg muscle volume (F(3, 43)=0.74, P=0.35). A significant group effect on peak isometric knee extension torque was found (F(3, 44)=3.06, P=0.038) with significantly lower values for group O vs. Y (adjusted P=0.0287). Kruskal-Wallis tests did not reveal a significant group effect on peak isometric knee flexion torque (H(3)=4.68, P=0.197) but did reveal significant group effects on peak isokinetic knee extension (H(3)=25.96, P<0.0001) and flexion (H(3)=25.98, P<0.0001) torque, with significantly greater peak isokinetic knee extension torque in group Y vs. O (adjusted P<0.0001), vs. IO (adjusted P=0.0024) and vs. TO (adjusted P=0.0178) and significantly greater peak isokinetic knee flexion torque in group Y vs. O (adjusted P<0.0001) and vs. IO (adjusted P=0.0022). Similar to the muscle fibre type characteristics, these data confirm a range of muscle strength values in our analysed participants. The muscle volume and strength data are displayed in the supplemental figures 1 and 2.

**Figure 2:**
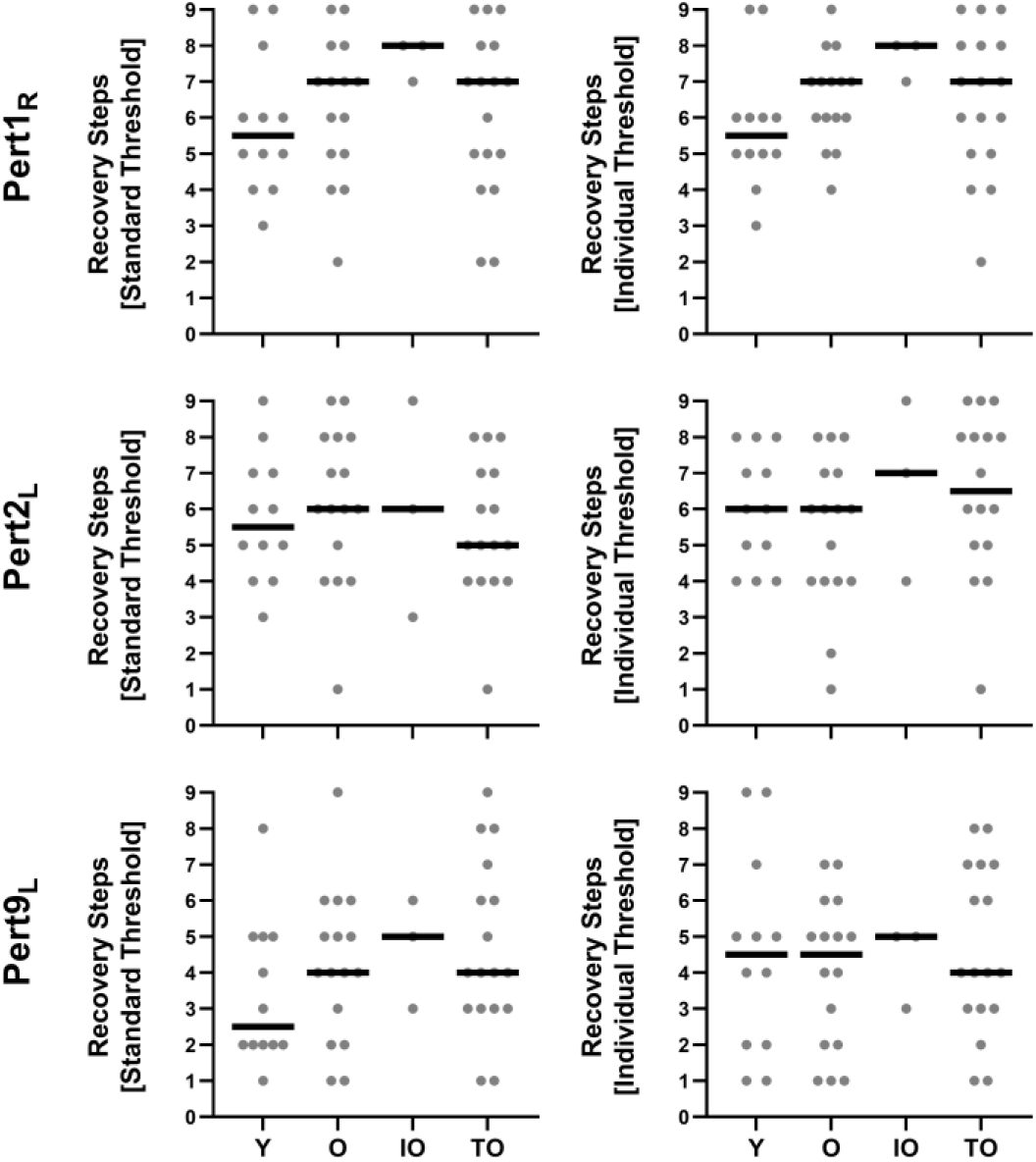
Group median and individual values for the number of recovery steps determined by the standard and individual thresholds for all the first perturbation to each leg (Pert1_R_ and Pert2_L_) and the final left leg perturbation (Pert9_L_) for young adults (Y), older adults with normal physical activity (O), trained older adults (TO) and physically impaired older adults (IO).

Kruskal-Wallis tests did not reveal a significant group effect on the number of recovery steps for any perturbation or recovery step threshold (Fig. 2; Standard Threshold: H(3)=2.35, P=0.5, H(3)=1.2, P=0.75 and H(3)=2.66, P=0.45 for Pert1_R_, Pert2_L_ and Pert9_L_, respectively; Individual Threshold: H(3)=4.6, P=0.2, H(3)=2.68, P=0.44 and H(3)=0.36, P=0.95 for Pert1_R_, Pert2_L_ and Pert9_L_, respectively). However, as can be seen in Figure 2, a large range of numbers of recovery steps was observed across the participants.

### No Associations between Type II Fibre Characteristics and Balance Recovery

To test our hypothesis of a relationship between Type II muscle fibre characteristics (size and proportion) and stability recovery (number of recovery steps needed following a novel perturbation), Spearman correlations were conducted between these outcomes. These revealed no significant associations between the number of recovery steps following Pert1_R_ and Pert2_L_ and the three Type II fibre characteristics: Type II fibre percentage, Type II fibre CSA and the percentage of total CSA taken up by Type II fibres (Table 2).

**Table 2:** Spearman Correlation Results for Recovery Steps and Type II Fibre Characteristics

|  |  | Pert1<sub>R</sub> |  | Pert2<sub>L</sub> |  | Pert9<sub>L</sub> |  |
| --- | --- | --- | --- | --- | --- | --- | --- |
|  |  | Standard Threshold | Individual Threshold | Standard Threshold | Individual Threshold | Standard Threshold | Individual Threshold |
| Type II Fibre Content [%] | *Spearman r* | 0.127 | 0.159 | 0.202 | -0.036 | 0.043 | -0.111 |
|  | *95% CI* | -0.175 to 0.407 | -0.143 to 0.434 | -0.100 to 0.469 | -0.327 to 0.262 | -0.252 to 0.331 | -0.391 to 0.187 |
|  | *P* | 0.395 | 0.286 | 0.174 | 0.812 | 0.771 | 0.452 |
| Type II Fibre Mean CSA [µm<sup>2</sup>] | *Spearman r* | -0.026 | -0.027 | 0.103 | 0.056 | -0.214 | -0.048 |
|  | *95% CI* | -0.319 to 0.271 | -0.320 to 0.270 | -0.198 to 0.387 | -0.243 to 0.345 | -0.476 to 0.083 | -0.336 to 0.247 |
|  | *P* | 0.861 | 0.856 | 0.490 | 0.710 | 0.144 | 0.745 |
| CSA % from Type II Fibres | *Spearman r* | 0.143 | 0.120 | 0.194 | -0.125 | -0.085 | -0.161 |
|  | *95% CI* | -0.159 to 0.420 | -0.182 to 0.401 | -0.107 to 0.463 | -0.405 to 0.177 | -0.367 to 0.213 | -0.433 to 0.137 |
|  | *P* | 0.338 | 0.422 | 0.191 | 0.404 | 0.568 | 0.273 |

To provide further context for the muscle fibre characteristic associations, associations between the number of recovery steps and upper leg muscle volume and torque were also analysed. These Spearman correlations revealed no significant between upper leg muscle volume and the number of recovery steps following Pert1_R_ and Pert2_L_ (Supplementary Table 1). They revealed no significant associations between peak isometric and isokinetic knee extension and flexion torque and the number of recovery steps following Pert1_R_ and one (of 8) moderate, significant association for Pert2_L_ (significant for Peak Isometric Knee Flexion Torque; Supplementary Table 1). Note that to provide further insight into these relationships, we also ran *post hoc* Spearman correlations between the fibre type characteristics and the muscle volume and torque outcomes, which revealed significant correlations between all pairs of muscle volume and torque outcomes, four (out of five) significant correlations between Type II fibre CSA and the muscle volume and torque outcomes but no significant correlations between Type II muscle fibre content or percentage of CSA from Type II fibres and the muscle volume and torque outcomes (Supplementary Table 2).

### No Associations between Type II Fibre Characteristics and Adapted Balance Recovery

To explore the potential role of Type II muscle fibre characteristics in adaptability of stability recovery responses, associations between the recovery steps after eight repeated perturbations to the left leg (Pert9_L_) and the Type II muscle fibre characteristics were conducted. Spearman correlations revealed no significant associations between the number of recovery steps following Pert9_L_ and the three Type II fibre characteristics: Type II fibre percentage, Type II fibre CSA and the percentage of total CSA taken up by Type II fibres (Table 2).

Again, to provide additional information, associations between the Pert9_L_ recovery steps and muscle volume and torque were also conducted. In general agreement with the fibre type correlations, Spearman correlations revealed no significant associations between upper leg muscle volume and the number of recovery steps following Pert9_L_, and only one (of 8) moderate, significant associations between peak isometric and isokinetic knee extension and flexion torque and the number of recovery steps following Pert9_L_ (Significant for Peak Isometric Knee Flexion Torque; Supplementary Table 1).

## Discussion

In this study, we aimed to provide the first data on whether or not Type II muscle fibre characteristics associate with reactive balance during walking in young and older adults with varying muscle fibre type composition. We hypothesised that there would be a small to moderate relationship between Type II muscle fibre characteristics (size and proportion) and stability recovery (number of recovery steps needed following a novel perturbation). We conducted correlation analyses on our data from 48 young and older participants, with varying physical activity levels, muscle fibre type characteristics, muscle volume and strength, as well as stability recovery performance, and were not able to confirm this hypothesis, as we did not observe such correlations. The results of our secondary, exploratory correlations with muscle volume and strength generally aligned with the fibre type characteristic results, since associations between these parameters and stability recovery were also virtually absent.

The current study found no significant differences in muscle fibre size in older adults compared to younger adults. Thus, we found no age-related differences between young and older participants who were not engaged in structured physical exercise but possessed comparable physical activity levels (~10K steps/day) well above the general recommendation [64]. These results appear in contrast to previous studies reporting a substantial decrease in muscle fibre size, especially in Type II fibres, with age [50,51,53]. However, most of these studies did not measure and control for an age-related decrease in physical activity [50,51,53], which could explain muscle fibre atrophy [59]. In line with our data, St-Jean-Pelletier et al. [52] observed an age-related decrease in muscle Type II fibre size in sedentary middle-aged and older adults (~7K vs. ~6.3K steps/day on average) but not in physically active middle-aged and older adults (~12.9K vs. ~12.6K steps/day). These results further support the hypothesis that muscle fibre atrophy is not simply a function of ageing *per se* but might also be driven by age-related physical inactivity and immobilization. However, it should be kept in mind that while the ANOVA was not significant, our *post hoc* analyses comparing the young adults with all older adults and with group O each found a significant effect of age on Type II fibre CSA (Hedges *g*=0.82 and *g*=0.84, respectively).

In the present study, the relative proportion of Type I muscle fibres was larger in endurance-trained older adults when compared to older adults with normal physical activity levels and young individuals. Although individual studies have shown conflicting findings regarding muscle fibre type distribution between young and older endurance-trained individuals [80,81], our results are in line with evidence indicating greater proportions of Type I fibres in endurance-trained older adults when compared to untrained older adults and untrained young individuals [57–59]. While evidence suggests long-term strength training provides a strong stimulus for the preservation of the structural and mechanical characteristics of skeletal muscle during ageing [54,82], it remains unclear to what extent aerobic training alone can counteract these age-related changes [80,83,84]. However, endurance athletes may benefit from other adaptations that play a role in stability. One previous study reported that in a mixed group of older and younger adults, those who were recreational runners performed better on a forward falling task than the non-active participants, despite similar muscle strength values [30]. The authors suggested that runners frequently manage the stability of their body position during running and may therefore be better able to perform such destabilising tasks [30]. In our study, of the 17 participants in group TO, eight participated in regular running and a further two participated in badminton and tennis, which also involve multidirectional stepping and change of direction, and these individuals may have compensated their reduction in Type II fibre size and number with relatively more refined stability control.

The literature is currently mixed regarding the role of muscle strength in balance and falls [9–13,17,39–42]. Despite this, small to moderate associations between lower limb muscle strength and balance recovery performance following laboratory-based perturbations [28–38] led us to our hypothesis of a small to moderate relationship between Type II muscle fibre properties and balance recovery performance from a sudden walking perturbation. Our results suggest that the muscle fibre properties themselves are not critical factors for reactive balance recovery during walking, due to the lack of correlations, the inconsistent direction of the (non-significant) correlations, as well as the fact that the trained older group did not require a significantly different number of recovery steps, despite a significantly different proportion of the muscle fibre CSA coming from Type II fibres. In order to check if the younger adults’ data were unduly influencing these results, we repeated the Spearmen correlation analyses *post hoc* with only the older participants included and these demonstrated similar results, with only one of the 18 associations being significant (Supplementary Table 3). Our collective muscle volume and strength correlation results were also mostly not statistically significant. These results may, to some extent, explain some of the mixed findings in the literature. Strength testing or training that primarily targets the muscle tissue properties and neglects other factors (such as motor coordination, response/movement speed, functionally relevant movements, among others; see Tieland et al. [47] and Hunter et al. [85] for reviews) may not capture critical elements of muscle function relevant for reactive balance control. That being said, the success of the recovery actions may not critically depend on maximal muscle strength at all. For example, one biomechanical modelling study found that, for backward balance loss, there is quite some overlap in the feasible regions for balance recovery during simulations with strength characteristics of young and older adults and recovery was only limited by strength during very short or very long recovery steps [86]. This is also supported by other studies demonstrating that older adults can improve their balance recovery responses to a level similar to young adults when repeatedly exposed to perturbations (acutely and over longer time periods), without changes in muscle strength [36,87]. As falls among community-dwelling older adults usually occur during dynamic movement [20–27], specificity of both strength and balance assessment and training should be key considerations in fall prevention.

### Study Limitations

Some limitations of the current study should be considered. Regarding the walking perturbation protocol, our participants were informed that they would complete a walking balance challenge but were unaware of the specifics of the perturbations and no warnings or cues were given during the trial. However, beneficial effects of increased awareness of perturbations on stability recovery performance following trips have been shown [88,89], even when the specifics are unknown. This may have resulted in better balance recovery performance than might be expected in natural settings. Regarding our outcome measure, the number of recovery steps, this is not a standard measure of balance recovery following perturbations to walking. However, no gold-standard outcome measure exists for this purpose. Other possibilities would be to use specific biomechanical variables such as the margin of stability or trunk lean at a specific time point. However, due to the heterogeneity in the balance recovery responses to such perturbations, using such specific measures at specific time points has limitations when trying to capture “overall” balance recovery during walking. We feel that the current measure gives a more general overall indicator that is more robust to individual differences in response behaviour. It should also be kept in mind that our muscle biopsies came from the vastus lateralis muscle which is not necessarily the most critical muscle for gait [2–4] (although it likely plays an important role in leg extension and energy absorption during the initial recovery steps) nor entirely representative of the fibre type properties from other lower limb muscles [90–92]. Finally, one potential limitation is the exploratory nature of some of the correlations and the lack of *a priori* correction for multiple comparisons. In general, despite the relatively high number of correlations performed, the lack of significance and the inconsistency in the direction of the correlations leads us to believe that our conclusions are justified and not dependent on spurious correlations (significant or not).

### Conclusions

The current results indicate that Type II muscle fibre properties (proportion and dimensions) are not significantly associated with reactive balance recovery following perturbations to walking in young and older adults. Upper leg muscle volume and knee flexion and extension torque also failed to show a significant relationship with reactive balance recovery. These results have implications for muscle strength testing and training in fall prevention, as it appears that the muscle tissue properties are not key factors for balance recovery.

## Acknowledgements

The authors thank Matthijs Hesselink for helpful discussions regarding the fibre type analysis, Tim Snijders for helpful comments and discussion on an earlier draft of the manuscript, Wouter Bijnens for assistance with the walking measurements, Paul Willems for technical support and Pascal Rense and Colin Dohmen for assistance with participant recruitment and various measurements.

## Author Contributions

CM: Conceptualisation, Data Curation, Formal Analysis, Investigation, Methodology, Visualisation, Writing – Original Draft, Writing – Reviewing & Editing.

LG: Conceptualisation, Data Curation, Funding Acquisition, Investigation, Project Administration, Writing – Original Draft, Writing – Reviewing & Editing.

GS: Investigation, Visualisation, Writing – Reviewing & Editing.

EKM: Investigation, Writing – Reviewing & Editing.

JAJ: Investigation, Writing – Reviewing & Editing.

AG: Investigation, Software, Writing – Reviewing & Editing.

KM: Conceptualisation, Resources, Supervision, Writing – Reviewing & Editing.

JH: Conceptualisation, Funding Acquisition, Project Administration, Resources, Supervision, Writing – Reviewing & Editing.

## Funding

CM was financially supported by the NUTRIM Graduate Programme of Maastricht University Medical Centre+. LG and JH are financially supported by the TIFN research program Mitochondrial Health (ALWTF.2015.5). The project is partly organised by and executed under the auspices of TiFN, a public-private partnership on precompetitive research in food and nutrition.

## Competing Interests

The authors declare no competing interests. Danone Nutricia Research, Friesland Campina, the Netherlands Organisation for Scientific Research and the Top-sector Agri&Food are sponsors of the TIFN program and partly financed the project. They had no role in data collection and analysis, and the decision to publish.

## Notes

### Competing Interest Statement

The authors have declared no competing interest.

