## Supplementary Material for "Type II muscle fibre properties are not associated with balance recovery following large perturbations during walking in young and older adults"


### Supplement Results

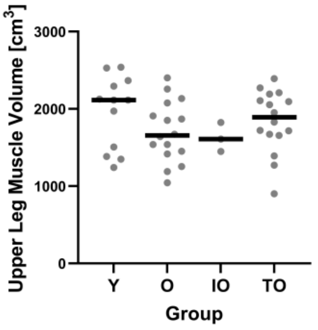


***Supplement Fig. 1**: Group median and individual upper leg muscle volume for young adults (Y), older adults with normal physical activity (O), trained older adults (TO) and physically impaired older adults (IO).*

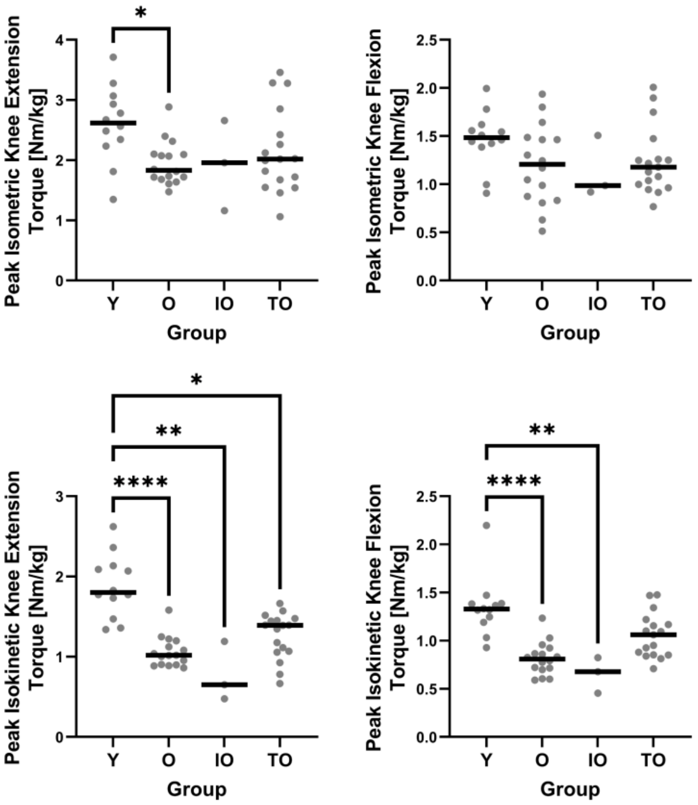


***Supplement Figure 2:** Group median and individual values for peak isometric and isokinetic knee extension and flexion torque for young adults (Y), older adults with normal physical activity (O), trained older adults (TO) and physically impaired older adults (IO). *: P<0.05 **P<0.01 ****:P<0.0001*

**Supplement Table 1:** Spearman correlation results for upper leg muscle volume, peak isometric and isokinetic knee extension and flexion torque and $Pert1_R$, $Pert2_L$ and $Pert9_L$

| | | **$Pert1_R$** | | **$Pert2_L$** | | **$Pert9_L$** | |
|---|---|---|---|---|---|---|---|
| | | *Standard Threshold* | *Individual Threshold* | *Standard Threshold* | *Individual Threshold* | *Standard Threshold* | *Individual Threshold* |
| **Upper Leg Muscle Volume** | *Spearman r* | -0.1226 | -0.01128 | -0.01661 | -0.1461 | -0.1766 | -0.1825 |
| | *95% CI* | -0.4061 to 0.1825 | -0.3086 to 0.2881 | -0.3134 to 0.2832 | -0.4259 to 0.1592 | -0.4484 to 0.1251 | -0.4532 to 0.1191 |
| | *P* | 0.417 | 0.9407 | 0.9127 | 0.3327 | 0.235 | 0.2195 |
| **Peak Isometric Knee Extension Torque** | *Spearman r* | 0.1023 | 0.09993 | -0.06967 | -0.2268 | -0.03795 | -0.1397 |
| | *95% CI* | -0.1989 to 0.3858 | -0.2012 to 0.3838 | -0.3575 to 0.2302 | -0.4892 to 0.07325 | -0.3264 to 0.2570 | -0.4148 to 0.1589 |
| | *P* | 0.4939 | 0.5039 | 0.6417 | 0.1252 | 0.7979 | 0.3437 |
| **Peak Isometric Knee Flexion Torque** | *Spearman r* | -0.01427 | 0.002994 | -0.2043 | -0.3262 | -0.2399 | -0.3902 |
| | *95% CI* | -0.3081 to 0.2821 | -0.2924 to 0.2979 | -0.4711 to 0.09667 | -0.5668 to -0.03440 | -0.4971 to 0.05612 | -0.6125 to -0.1108 |
| | *P* | 0.9242 | 0.9841 | 0.1683 | **0.0252** | 0.1006 | **0.0061** |
| **Peak Isokinetic Knee Extension Torque** | *Spearman r* | -0.1622 | -0.2081 | -0.2041 | -0.06516 | -0.1998 | -0.1164 |
| | *95% CI* | -0.4365 to 0.1396 | -0.4741 to 0.09277 | -0.4709 to 0.09691 | -0.3535 to 0.2345 | -0.4647 to 0.09799 | -0.3950 to 0.1819 |
| | *P* | 0.276 | 0.1604 | 0.1688 | 0.6635 | 0.1734 | 0.4308 |
| **Peak Isokinetic Knee Flexion Torque** | *Spearman r* | -0.08 | -0.1216 | -0.2476 | -0.07478 | -0.1597 | -0.07469 |
| | *95% CI* | -0.3665 to 0.2204 | -0.4023 to 0.1800 | -0.5058 to 0.05127 | -0.3620 to 0.2254 | -0.4317 to 0.1388 | -0.3589 to 0.2222 |
| | *P* | 0.593 | 0.4155 | 0.0933 | 0.6174 | 0.2781 | 0.6139 |

**Supplement Table 2:** Spearman correlation matrix for upper leg muscle volume, peak isometric and isokinetic knee extension and flexion torque, Type II fibre content, Type II fibre mean CSA and CSA percentage from Type II fibres

| | | **Upper Leg Muscle Volume** | **Peak Isometric Knee Extension Torque** | **Peak Isometric Knee Flexion Torque** | **Peak Isokinetic Knee Extension Torque** | **Peak Isokinetic Knee Flexion Torque** | **Type II Fibre Content [%]** | **Type II Fibre Mean CSA [µm$^2$]** | **CSA % from Type II Fibres** |
|---|---|---|---|---|---|---|---|---|---|
| **Upper Leg Muscle Volume** | *Spearman r* | | | | | | | | |
| | *95% CI* | | | | | | | | |
| | *P* | | | | | | | | |
| **Peak Isometric Knee Extension Torque** | *Spearman r* | 0.405412 | | | | | | | |
| | *95% CI* | 0.1252 to 0.6257 | | | | | | | |
| | *P* | **0.0047** | | | | | | | |
| **Peak Isometric Knee Flexion Torque** | *Spearman r* | 0.533534 | 0.646 | | | | | | |
| | *95% CI* | 0.2829 to 0.7159 | 0.4361 to 0.7891 | | | | | | |
| | *P* | **0.0001** | **<0.0001** | | | | | | |
| **Peak Isokinetic Knee Extension Torque** | *Spearman r* | 0.631707 | 0.608 | 0.528 | | | | | |
| | *95% CI* | 0.4137 to 0.7812 | 0.3839 to 0.7642 | 0.2790 to 0.7105 | | | | | |
| | *P* | **<0.0001** | **<0.0001** | **0.0001** | | | | | |
| **Peak Isokinetic Knee Flexion Torque** | *Spearman r* | 0.490634 | 0.670 | 0.538 | 0.877 | | | | |
| | *95% CI* | 0.2286 to 0.6864 | 0.4697 to 0.8045 | 0.2914 to 0.7171 | 0.7858 to 0.9305 | | | | |
| | *P* | **0.0005** | **<0.0001** | **0.0001** | **<0.0001** | | | | |
| **Type II Fibre Content [%]** | *Spearman r* | -0.04845 | 0.043 | 0.175 | 0.045 | -0.118 | | | |
| | *95% CI* | -0.3388 to 0.2503 | -0.2523 to 0.3308 | -0.1234 to 0.4443 | -0.2508 to 0.3322 | -0.3961 to 0.1806 | | | |
| | *P* | 0.7464 | 0.7723 | 0.2342 | 0.7639 | 0.4257 | | | |
| **Type II Fibre Mean CSA [µm$^2$]** | *Spearman r* | 0.570421 | 0.282 | 0.297 | 0.564 | 0.405 | -0.018 | | |
| | *95% CI* | 0.3310 to 0.7408 | -0.01117 to 0.5302 | 0.005062 to 0.5418 | 0.3256 to 0.7350 | 0.1284 to 0.6235 | -0.2946 to 0.2610 | | |
| | *P* | **<0.0001** | 0.0523 | **0.0406** | **<0.0001** | **0.0043** | 0.8970 | | |
| **CSA % from Type II Fibres** | *Spearman r* | 0.066374 | 0.187 | 0.297 | 0.185 | 0.048 | 0.855 | 0.331 | |
| | *95% CI* | -0.2334 to 0.3546 | -0.1114 to 0.4540 | 0.005538 to 0.5421 | -0.1132 to 0.4526 | -0.2473 to 0.3355 | 0.7566 to 0.9153 | 0.05832 to 0.5575 | |
| | *P* | 0.6576 | 0.2038 | **0.0403** | 0.2082 | 0.7449 | **<0.0001** | **0.0155** | |

**Supplementary Table 3**: Spearman Correlation Results for Recovery Steps and Type II Fibre Characteristics in the Older Participants.

| | | **$Pert1_R$** | | **$Pert2_L$** | | **$Pert9_L$** | |
|---|---|---|---|---|---|---|---|
| | | *Standard Threshold* | *Individual Threshold* | *Standard Threshold* | *Individual Threshold* | *Standard Threshold* | *Individual Threshold* |
| **Type II Fibre Content [%]** | *Spearman r* | 0.2188 | 0.3559 | 0.1725 | -0.1154 | -0.08924 | -0.2365 |
| | *95% CI* | -0.1335 to 0.5220 | 0.01544 to 0.6224 | -0.1805 to 0.4861 | -0.4404 to 0.2362 | -0.4143 to 0.2560 | -0.5315 to 0.1098 |
| | *P* | 0.2066 | **0.0359** | 0.3217 | 0.5090 | 0.6048 | 0.1650 |
| **Type II Fibre Mean CSA [$\mu m^2$]** | *Spearman r* | 0.008247 | 0.001569 | 0.06739 | -0.009371 | -0.2108 | -0.08795 |
| | *95% CI* | -0.3350 to 0.3496 | -0.3409 to 0.3437 | -0.2814 to 0.4005 | -0.3506 to 0.3340 | -0.5119 to 0.1364 | -0.4132 to 0.2572 |
| | *P* | 0.9625 | 0.9929 | 0.7005 | 0.9574 | 0.2172 | 0.6100 |
| **CSA % from Type II Fibres** | *Spearman r* | 0.2514 | 0.3135 | 0.1707 | -0.2258 | -0.1783 | -0.3070 |
| | *95% CI* | -0.09948 to 0.5467 | -0.03227 to 0.5923 | -0.1823 to 0.4847 | -0.5273 to 0.1263 | -0.4866 to 0.1693 | -0.5840 to 0.03403 |
| | *P* | 0.1452 | 0.0667 | 0.3269 | 0.1922 | 0.2980 | 0.0686 |
